# Biochar effect - mediated ammonium transporter genes and GDH may be the core of improvement nitrogen assimilation in cotton seedling

**DOI:** 10.1101/2020.02.22.960997

**Authors:** Lei Feng, Guangmu Tang, Wanli Xu, Meiying Gu, Zengchao Geng

## Abstract

Biochar enhancement of nitrogen efficiency in crops is highly essential not only to reduce costs of agricultural production but also to conserve resources, lower energy consumption for products of these fertilizers, strengthen soil health, and eventually helps in slowing climate change; however nitrogen efficiency physiology by biochar effects is not clear. Here, we reported on the morphological, nitrogen metabolism and cytokinin, at seedling stage, under different layers of biochar and limited urea conditions grown in soil culture. Expression profile of miRNAs and AOB was further studied in fine and medium roots. It showed active root absorption area, fresh weight, and nitrogen agronomic efficiency responded significantly under biochar and reduction by 20% urea condition in the surface soil layer. Also, NR and GPT activity in fine roots remarkably increased with cytokinin, but decreased significantly in medium roots, meanwhile both NR and GDH activity did so. GOGAT activity was to be dependent with biochar and urea locations. In addition, *AMT1;1, gdh3* and *gdh2* in fine roots showed their up-regulation with reduction 20% urea and biochar. It revealed that co-expression of *gdh3* and *gdh2* in fine roots significantly affected nitrogen assimilation under reduction 20% urea with biochar on surface soil at seedling stage.

**Highlights:** The co-expression of ammonium transporter gene and *GDH* induced by biochar effect improves nitrogen efficiency and seedling growth.

These data emphasizes the importance of effects of cytokinin on nitrate reductase activity closely related to the position under biochar condition, which is a key element of enhancement nitrogen assimilation efficiency in cotton seedling.

Biochar addition applied into 0 to 10cm soil had a more positive effect on seedling growth than that into 10 to 20cm soil layers.

## 1 Introduce

Improving the nitrogen efficiency of crops can not only reduce the cost of planting, but also reduce the energy consumption caused by chemical fertilizers, thus fundamentally mitigating global climate change (Sinha et al., 2015). So regulation of nitrogen metabolism from molecular and physiological level is a key link (Asif et al. 2020).

Biochar can significantly improve the nitrogen-urea utilization efficiency and crop productivity (Jeffery et al. 2011; Pandey et al. 2016). Studies supporting this view suggest that biochar increases rhizosphere microbial community diversity (Dilfuza et al. 2016; Kolton et al. 2017) or microbial biomass (Lu et al.2018), especially increasing the abundance of ammonia-oxidizing bacteria closely related to the nitrogen cycle (Wu et al. 2016), which in turn significantly increases the availability of nitrogen in the rhizosphere of crops (Song et al. 2014), resulting in increased nitrogen efficiency (Backer et al. 2017). Changes in biochar application patterns also may be a major cause, which are directly related to the effects of biochar. Besides, studies suggest that biochar induced root physiological and molecular changes may be the core of improving nitrogen efficiency (Farhangi-abriz & Torabian 2017; Noguera et al. 2012). Only Viger et al. (2014) reviewed that biochar effects regulate plant genes under abiotic stress, it seemed as biochar interfered with microbial signals (Masiello et al., 2013). However, the mechanism of root nitrogen metabolism involving biochar mediation has not yet been elucidated (Polzella et al. 2018).

The physiological and molecular variation of root or plant induced by biochar effect was strongly correlated with nitrogen metabolism (Backer et al. 2017), and its effect on physiological level of fine roots was even more significant (Razaq et al. 2017). From the study the authors found that biochar decreased ethylene concentration and increased the number of roots of tissue cultured poplar (Lonardo et al. 2013), which indicated that root characteristics were significantly affected by biochar. A recent research reported that 50 and 100 g kg^-1^ biochar have the same effect in improving nitrogen metabolism; it could be independent of biochar dose to a certain extent (Farhangi-abriz & Torabian 2018). In depth examination indicated that DOMs in biochar have promoted nitrogen assimilation and improved nitrogen efficiency by stimulating nitrate reductase and glutamine synthase gene expression (Bian et al. 2019). Another study demonstrated biochar promotes turnover of leaf proteases and thus increases rice biomass (Noguera et al. 2012). Hashem et al. (2019) further revealed that biochar enhanced the nitrogen assimilation efficiency of chickpeas by increasing nitrate reductase activity. There are more reports on enzymes in crop species and different organs, but interspecies’ root variation and nitrogen metabolism with biochar are not well understood.

The effects of biochar on hormone levels and spatial distribution patterns may be major factors that interfere with the activities of key enzymes involved in nitrogen metabolism (Spokas et al. 2010; Mehari et al., 2015; Farhangi-abriz & Torabian 2017). Waqas et al. (2014) ever considered that the changes of jasmonic acid signal reflected the alleviating effect of biochar on abiotic stress. Recent studies showed that biochar stimulated the gibberellins pathway and promoted the growth of tomato (*Solanum lycopersicum*) to some extent (French & iyer-pascuzzi 2018). On the contrary, Hale et al. (2015) considered that 600°C pyrolysis of pine sawdust biochar has no effect on auxin synthesis. These suggested that biochar may impact on plant endogenous hormones and depend on the type. Thusly, whether biochar can interfere with cytokinin metabolism and further affect *GDH* coding gene is not clear.

Although some studies have pointed out that *GDH* enzyme has not been detected in each organ compared with the wild type with *gdh1-2-3* mutant, and only *gdh2* mutant can reduce root *GDH* enzyme activity by 25%. Only *gdh3* mutants increase by 30% were present in the root system (Fontaine et al. 2012; Konishi et al. 2014). It can be inferred from that biochar may affect the synergistic effect of *AMT* and *GDH* genes, However, there is still insufficient evidence that biochar acted on nitrogen metabolism by affecting this metabolic pathway.

The objectives of this study were to evaluate changes in soil parameters (NH_4_^+^, NO_3_^-^, and AOB counts), nitrogen-metabolizing enzymes (nitrate reductase, glutamate dehydrogenase, glutamic-pyruvic transaminase, glutamic synthase), and biological character (root active absorbing area, biomass, nitrogen agronomic efficiency) caused by Bc addition with reduction of urea, meanwhile, to detect cotton AMT and *gdh* affectation on nitrogen metabolism induced by Bc; to determine how AMT and *gdh* expression discrepancy or consistency under ammonium assimilation pathway; and to explain how cytokinin stimulated nitrate metabolism contribute to the seedling growth. Three hypotheses were tested: (i) Bc addition alters soil bio-chemical properties, acting positively on soil nitrogen status and increasing ammonia-oxidizing bacteria number; (ii) these Bc-induced effects favor root intraspecific variation caused crosstalk between AMT1;1 and *gdh2*; and (iii) as a result, seedling in Bc with reduction of urea treated soil show a higher nitrogen metabolism and growth rate.

## 2 Material and Methods

### 2.1 Plant Growth Conditions

*Gossypium hirsutum* seeds were surface sterilized for 20min in sterile water, and placed on Petri dishes. After 48 to 72 h of incubation at 4°C in the dark, the plates were transferred to a growth pot in greenhouse. Seedlings were removed from the plates and transferred to soil culture units consisting of plastic containers containing 7.5kg gray desert soil and urea as sole N source. Each pot unit contained 12 plants. A pot experiment with five replications was conducted on April 5, 2019, in a glass greenhouse with a factorial design based on the randomized complete block design. Three doses of urea (i.e., 3.76 and 2.82 and 1.88 g°kg^-1^) chosen according to the conventional use of nitrogen fertilizer in Xinjiang Uygur Autonomous Region of cotton China and two biochar treatments (i.e., non-biochar and 37.28 g kg^-1^ soil) were used to test the cotton (*Gossypium hirsutum* 49).

The soil was mixed with biochar and distributed at 0~10 and 10~20cm layers into pots 20 cm in radius and 25 cm in height in amounts of 7.5 kg per pot. The properties of experimental soil present pH8.35, conductivity 0.22ms cm^-1^, organic carbon 6.91g kg^-1^, total nitrogen, total phosphorus and total potassium were 0.47, 0.59 and 16.35g kg^-1^, respectively. The pots were kept in controlled conditions in a glass greenhouse with a day and night temperature cycle of 30 and 5 °C, respectively, as well as 55~60% relative humidity, 150 W m^-2^ light intensity, and a 13 h photoperiod. Plants were irrigated every day with tap water in an amount comparable to field water capacity. Plants were grown for 40 d on a pot. Fine and medium roots were harvested 2 h for the metabolome and transcriptome analyses.

### 2.2 Determination of soil nitrate and ammonium nitrogen

Fresh soil samples were extracted by KCl solution and ammonium nitrogen and nitrate nitrogen in filtrate were determined by continuous flow analyzer.

### 2.3 Ammonia oxidizing bacteria

The 16s rDNA fragment of AOB was amplified by nested PCR (nest-PCR), and the primer sequence was used. F27/R1492 was the common primer for bacteria. CTO189F/CTO654R is a specific primer for AOB; F341/R518 is a V3 region specific primer for 16s rDNA.

AOB primers CTO189F - 5 ‘GCAGRAAAGYAGGGGATCG; CTO654R - 5 ‘CTAGGYTTGTAGTTTCAAACGC.

### 2.4 Enzyme Assays Analysis

For protein extraction, roots of 35-days-old Arabidopsis plants were harvested. Proteins were extracted from frozen root material stored at −80°C. All extractions were performed at 4°C. NADH-GDH activity was measured as described by Turano et al. (1996). Results for the NADH-GDH activities are presented as mean values for five plants with standard errors.

### 2.5 qRT-PCR

Total RNA was extracted from leaves and roots using TRIzol Reagent (Invitrogen) according to the manufacturer’s protocol. Genomic DNA was removed from the total RNA by treatment with amplification-grade DNaseI (Sigma-Aldrich) according to the manufacturer’s protocol.

### 2.6 Statistical Analysis

Genes with a Bonferroni *P* value ≤0.05 were considered as being differentially expressed, as described by Sarasin & Kauffmann (2008). In addition to the paired t test approach, the rank product method was used to detect differentially expressed genes according to different levels of FDR (Hong et al., 2006). We have chosen to present transcripts satisfying both the above family-wise error rate level and a FDR <0.0001% for an optimal interpretation of the transcriptome. Sorting genes by descending rank product values provided a hierarchical list based on both strength and reproducibility, which was used as an input to identify groups of genes with the same or related annotated function (Hong et al., 2006).

## 3 Results

### 3.1 Effects of nitrogen reduction combined with biochar on key enzymes of nitrogen metabolism

Nitrogen reduction combined with biochar strongly interfered with nitrogen metabolism of seedling roots, and there was a significant difference in the fluctuation of key enzymes of nitrogen metabolism. The GDH activity of root was still high when nitrogen application was reduced by 20% and 22.5g kg^-1^ biochar was optimum, among which the GDH activity of root <0.1mm treated by soaf increased by 2-folds. The GDH activity of roots in classification from 0.1 to 2mm universally decreased, with a maximum decrease of 91%. In addition, with only 50% nitrogen reduction, the GDH activity was all significantly reduced (*P*<0.05) (Figure 1A).

**Figure 1.**
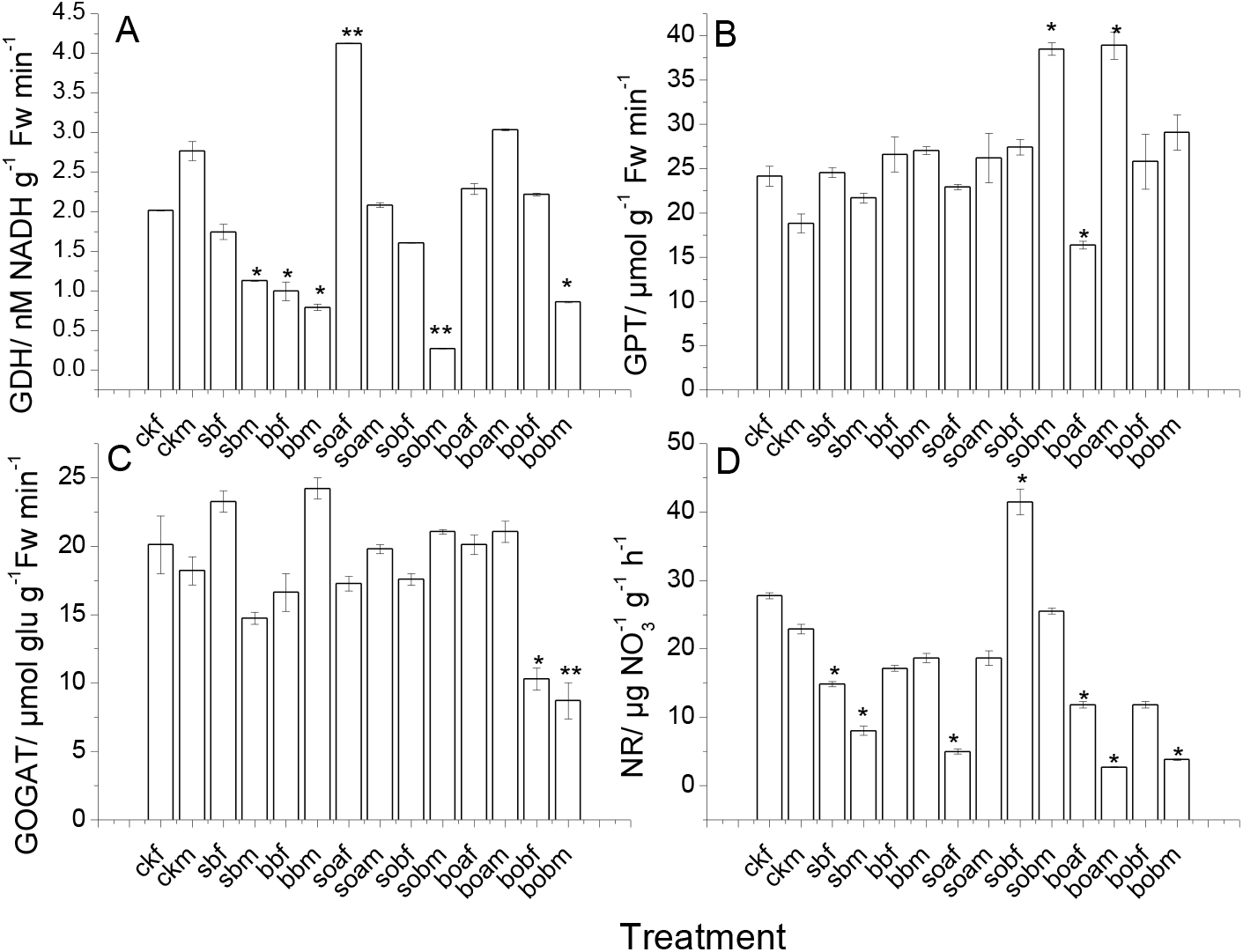
Changes of biochar applied to four key enzymes of nitrogen metabolism with classification of root less than 0.1mm (fine root) and between 0.1 and 2mm(medium root) under nitrogen reduction conditions. GDH (Glutamic dehydrogenase), GPT (Glutamic-pyruvic transaminase), GOGAT (Glutamic synthase), NR (Nitrate Reductase). There are seven treatments ck (control), sb (conventional 3.76 g nitrogen kg^-1^ soil fertilization in 0~10cm soil layers), bb (conventional nitrogen fertilization in 10~20cm soil layers), soa (Nitrogen was reduced by 20% on the basis of sb with 37.28 g biochar kg^-1^ soil), sob (Nitrogen was reduced by 50% on the basis of sb with 37.28 g biochar kg^-1^ soil), boa (Nitrogen was reduced by 20% on the basis of bb with 37.28 g biochar kg^-1^ soil), and bob (Nitrogen was reduced by 50% on the basis of bb with 37.28 g biochar kg-1 soil), respectively. The last letter “f” and “m” of each treatment indicates fine root and medium root separately. Results are presented as mean values for five plants with SE. Asterisk indicate significant differences with a confidence interval at \**P* < 0.05 and \*\**P*<0.01, respectively.

Nitrogen reduction combined with biochar did not significantly change the GPT activity, only sobm treatment significantly increased the GPT activity (*P*<0.05), boaf treatment significantly decreased (*P* <0.05), indicating that seedling medium roots may be more sensitive to nitrogen reduction with biochar effects. Since GPT can represent the geotropism of crop roots to a certain extent, medium roots have a higher probability of spreading to the deep, so the activity is relatively high (Figure 1B). The enzymatic activity of biochar GOGAT in combination with nitrogen reduction showed two trends. Second, reduction 50% nitrogen, applied to 10-20cm soil layer, fine roots (*P*<0.05) and medium roots (*P*<0.01) showed significant or extremely significant reduction (Figure 1C). For NR activity, except sobf treatment, all other treatments decreased to different degrees. Soaf treatment decreased by 83 %(Figure 1D).

Previous studies have shown that the transcription level is more significant in the regulation of nitrogen metabolism, and the ammonium absorption and transport gene (AMT) is mainly expressed in the root system, which is an important parameter for screening nitrogen-efficient varieties. As shown in Figure 2, The variation pattern of *AMT1;1* and *AMT1;3* gene expression level was similar in the whole roots, except that the expression level of medium root in soa treatment increased by 1.75-folds and that of fine root in Sob treatment increased by 78%, and the other treatments decreased with the reduction of nitrogen dose in biochar. Application of biochar at 0 to10cm soil layers was more conducive to stimulating root AMT expression, while application at 10~20cm soil layers had a stronger inhibitory effect (Figure 2A). Except for a 62% increase in the expression of *gdh2* in soaf treated (*d*<0.1mm) roots (hereinafter referred to as fine root), bbf, boaf and bobf treated fine roots decreased significantly, while sbf and sobf treated fine roots did not change significantly. Bob treatment (0.1mm<d<2mm) root (hereinafter referred to as the medium root) significantly decreased, while the rest of the treatment did not change significantly (Figure 2B).

**Figure 2.**
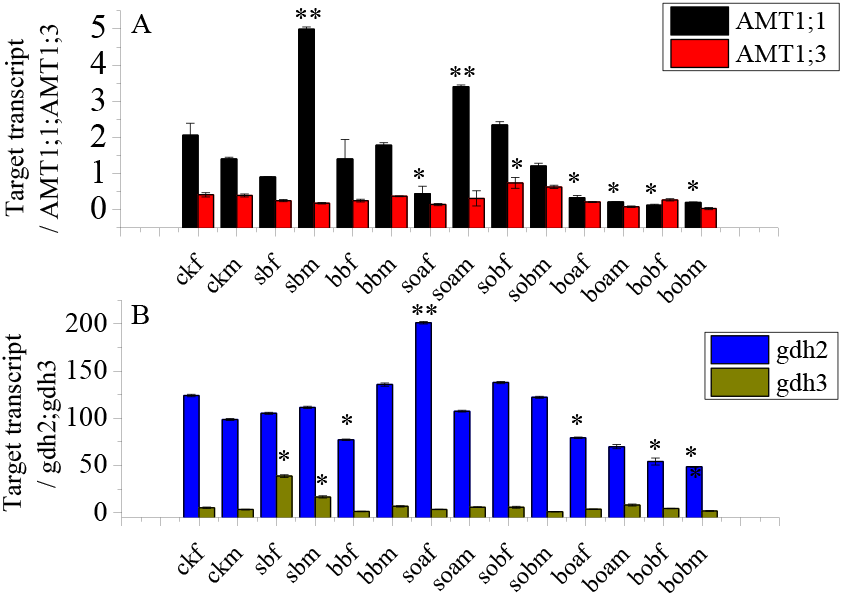
Quantification of transcript abundance in fine root and medium root of *Gossypium hirsutum* for the Genes Encoding *AMT* and *gdh*. qRT-PCR reactions were performed on total RNA extracted from the fine root and main root under reduction and biochar condition shown above. The level of expression of the AMT1;1, AMT1;3, gdh2, and gdh3genes was quantified for each of the 14 treatments. Results are presented as mean values for five plants with SE. Asterisk indicate significant differences with a confidence interval at \**P* < 0.05 and \*\**P*<0.01, respectively.

### 3.2 Crosstalk between cytokinin and nitrate reductase

The activity of NR was closely related to the location where it was distributed. CTK in fine root significantly promoted the increase of NR activity (Figure 3A), while in medium roots, NR activity significantly decreased with the growth of CTK activity (Figure 3B). Moreover, the activity range of fine roots was more extensive, ranging from 9 to 45, and that of medium root ranged from 16 to 27. Similarly, variance analysis showed that biochar effect increased the synergistic effect between CTK and NR, which could be one of the key reasons for biochar to enhance nitrogen metabolism.

**Figure 3.**
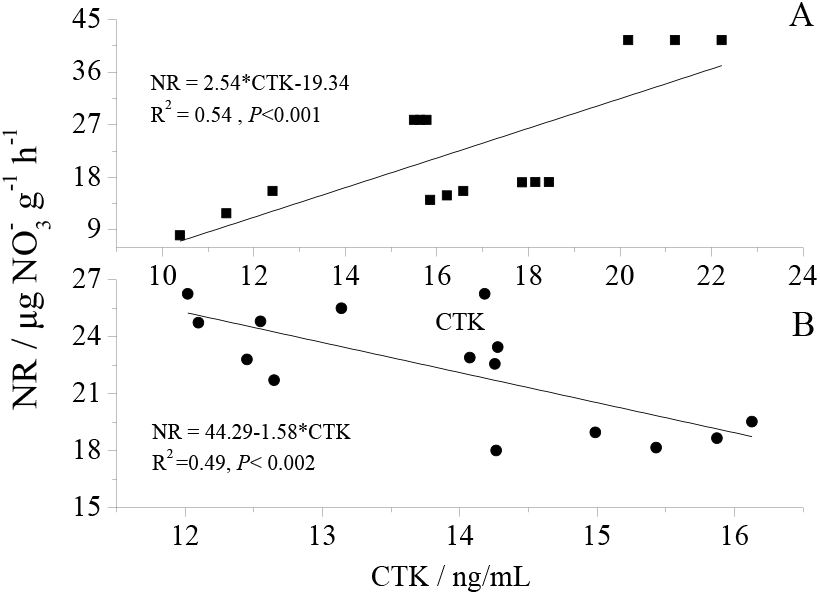
The activation and inhibition of cytokinin on nitrate reductase activity in root system. A and B represents the action of endogenous hormones and enzymes in fine roots and medium root, respectively.

### 3.3 Changes of nitrate and ammonium nitrogen in rhizosphere

The contents of nitrate nitrogen in different soil layers had complex regularity in the rhizosphere of different root sequences. The content of nitrate nitrogen in rhizosphere was the highest in both deep and shallow application. The content of nitrate nitrogen in the rhizosphere of 0~10cm was slightly higher than that of the fine roots. The nitrogen-nitrogen content in the rhizosphere of biochar with 20% nitrogen reduction was significantly higher than that of the surface application except conventional application. The difference between 10~20cm fine roots and medium roots was large, among which boa treated fine roots were 42% medium roots, and Bob treated fine roots had a maximum content of 6.33.This may indicate that nutrient uptake and utilization of crop roots in arid regions are insufficient by using <2mm thin roots, and further dividing thin roots into < 0.1mm and 0.1 < *d* < 2mm can describe the changes of available nitrogen content in rhizosphere relatively accurately (table 1).

**Table 1.** Excitation effect of biochar on nitrogen (NO_3_^-^, NH_4_^+^) in different soil layers and rhizosphere

| $\text{NO}_3^-$ concentration | | | | |
| --- | --- | --- | --- | --- |
|  | 0~10cm |  | 10~20cm |  |
| Treatment | Fine root | Medium root | Fine root | Medium root |
| ck | 0.13(0.04)d | 0.21(0.11)d | 0.11(0.01)d | 0.42(0.03)d |
| s | 7.33(1.02)a | 8.22(0.58)a | 3.46(0.33)bc | 4.62(0.29)c |
| b | 3.28(0.31)c | 3.75(1.04)c | 1.04(0.19)cd | 1.09(0.38)d |
| soa | 7.65(0.82)a | 9.58(0.54)a | 4.12(0.08)b | 9.77(0.36)a |
| sob | 5.62(1.12)b | 6.33(0.24)b | 3.09(0.25)bc | 4.73(0.17)c |
| boa | 5.09(0.49)bc | 5.89(0.78)b | 3.01(0.37)bc | 7.09(0.28)b |
| bob | 4.16(0.08)bc | 4.57(0.04)bc | 6.33(0.19)a | 9.69(0.05)a |
| $\text{NH}_4^+$ concentration | | | | |
|  | 0~10cm |  | 10~20cm |  |
| Treatment | Fine root | Medium root | Fine root | Medium root |
| ck | 0.076(0.021)bc | 0.083(0.009)b | 0.084(0.013)b | 0.088(0.014)b |
| s | 0.061(0.005)c | 0.069(0.003)c | 0.041(0.009)c | 0.042(0.017)c |
| b | 0.089(0.009)b | 0.095(0.013)ab | 0.092(0.016)ab | 0.099(0.006)ab |
| soa | 0.104(0.033)a | 0.127(0.012)a | 0.107(0.029)a | 0.101(0.024)ab |
| sob | 0.092(0.005)ab | 0.086(0.015)b | 0.085(0.008)b | 0.089(0.019)b |
| boa | 0.048(0.008)d | 0.054(0.002)c | 0.117(0.028)a | 0.125(0.003)a |
| bob | 0.083(0.021)b | 0.076(0.004)bc | 0.104(0.002)a | 0.118(0.002)a |

Note: The different lowercase letter indicate significant differences with a confidence interval at $*P < 0.05$ .

The regularity of ammonium nitrogen is similar to that of nitrate nitrogen. The contents of fine roots and medium rhizosphere with 20% nitrogen reduction were significantly higher than ck. Compared with the 10~20cm fine roots, the 0~10cm soil layer showed a decrease of 0.003mg/L, while the medium roots increased by 0.026mg/L. When applied at 10~20cm, boa treated fine root and medium root ammonium nitrogen at 0~10cm reduced by 59% and 57%, respectively (Table 2).

**Table 2.** AOB copy number at root class

| Treatment | Fine root | Medium root |
| --- | --- | --- |
| ck | $4.71 \times 10^7$ | $5.21 \times 10^7$ |
| s | $8.65 \times 10^7$ | $4.98 \times 10^7$ |
| b | $6.86 \times 10^7$ | $4.53 \times 10^7$ |
| soa | $7.29 \times 10^7$ | $9.70 \times 10^7$ |
| sob | $6.58 \times 10^7$ | $9.28 \times 10^7$ |
| boa | $5.44 \times 10^7$ | $1.08 \times 10^8$ |
| bob | $7.83 \times 10^7$ | $6.95 \times 10^7$ |

Note: Fine root less than 0.1mm and medium root from 0.1 to 2mm.

### 3.4 Ammonia-oxidizing bacteria copy number

The number of AOB copies in fine and medium rhizosphere was between 4.71~8.65×10^7^ and 4.53~10.8×10^7^, respectively, indicating that the rhizosphere of roots between 0.1~2mm provided relatively sufficient nitrogen sources for AOB flora. The highest AOB copy number was 8.65×107, which was not significantly different from soa, SOB and Bob treatment, but significantly higher than boa treatment. The amount of AOB in the medium rhizosphere decreased with conventional fertilizer application and soil layers of 0~10cm and 10~20cm.The amount of AOB copies increased by 1.86 and 1.78-folds respectively. When 20% and 50% nitrogen reduction were applied to soil layer 0~10cm and biochar were applied. The amount of AOB in the medium rhizosphere treated by boa reached 1.08×10^8^ (Table 3).

**Table 3.**
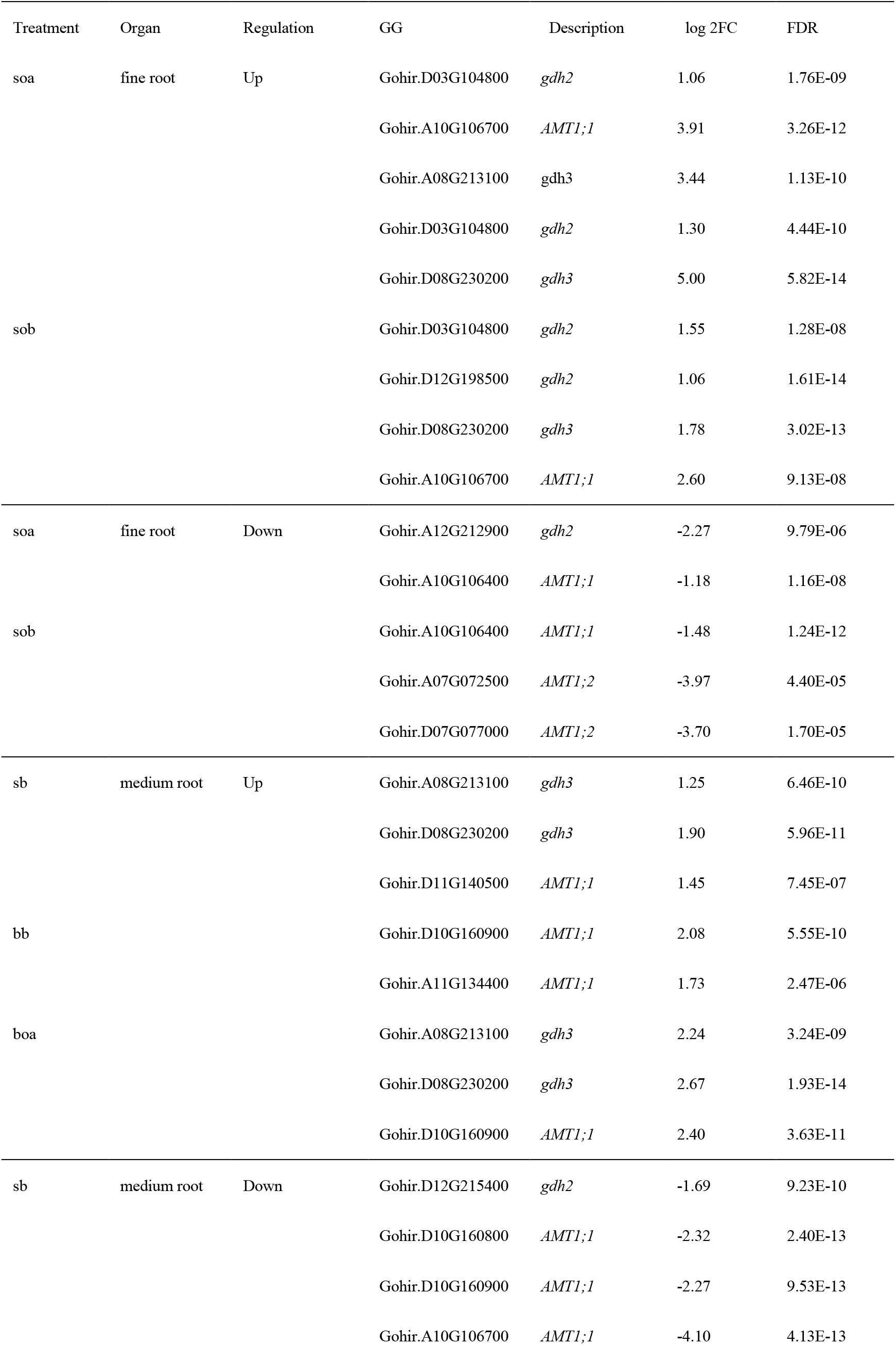
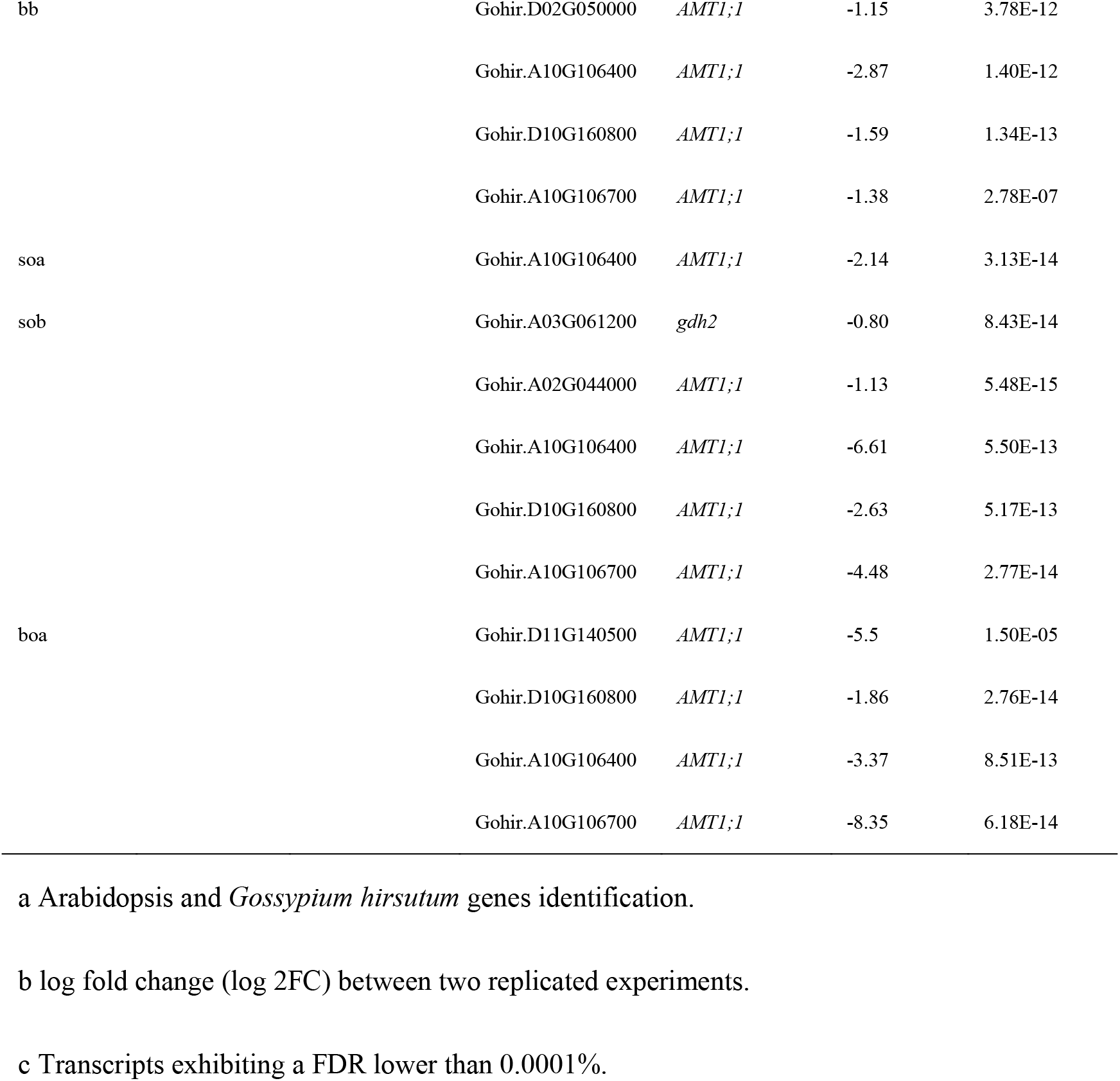
Transcript Abundance That Is Significantly Altered in the *Gossypium hirsutum gdh* and AMT under biochar and urea condition

| Treatment | Organ | Regulation | GG | Description | log 2FC | FDR |
| --- | --- | --- | --- | --- | --- | --- |
| soa | fine root | Up | Gohir.D03G104800 | *gdh2* | 1.06 | 1.76E-09 |
|  |  |  | Gohir.A10G106700 | *AMT1;1* | 3.91 | 3.26E-12 |
|  |  |  | Gohir.A08G213100 | *gdh3* | 3.44 | 1.13E-10 |
|  |  |  | Gohir.D03G104800 | *gdh2* | 1.30 | 4.44E-10 |
|  |  |  | Gohir.D08G230200 | *gdh3* | 5.00 | 5.82E-14 |
| sob |  |  | Gohir.D03G104800 | *gdh2* | 1.55 | 1.28E-08 |
|  |  |  | Gohir.D12G198500 | *gdh2* | 1.06 | 1.61E-14 |
|  |  |  | Gohir.D08G230200 | *gdh3* | 1.78 | 3.02E-13 |
|  |  |  | Gohir.A10G106700 | *AMT1;1* | 2.60 | 9.13E-08 |
| soa | fine root | Down | Gohir.A12G212900 | *gdh2* | -2.27 | 9.79E-06 |
|  |  |  | Gohir.A10G106400 | *AMT1;1* | -1.18 | 1.16E-08 |
| sob |  |  | Gohir.A10G106400 | *AMT1;1* | -1.48 | 1.24E-12 |
|  |  |  | Gohir.A07G072500 | *AMT1;2* | -3.97 | 4.40E-05 |
|  |  |  | Gohir.D07G077000 | *AMT1;2* | -3.70 | 1.70E-05 |
| sb | medium root | Up | Gohir.A08G213100 | *gdh3* | 1.25 | 6.46E-10 |
|  |  |  | Gohir.D08G230200 | *gdh3* | 1.90 | 5.96E-11 |
|  |  |  | Gohir.D11G140500 | *AMT1;1* | 1.45 | 7.45E-07 |
| bb |  |  | Gohir.D10G160900 | *AMT1;1* | 2.08 | 5.55E-10 |
|  |  |  | Gohir.A11G134400 | *AMT1;1* | 1.73 | 2.47E-06 |
| boa |  |  | Gohir.A08G213100 | *gdh3* | 2.24 | 3.24E-09 |
|  |  |  | Gohir.D08G230200 | *gdh3* | 2.67 | 1.93E-14 |
|  |  |  | Gohir.D10G160900 | *AMT1;1* | 2.40 | 3.63E-11 |
| sb | medium root | Down | Gohir.D12G215400 | *gdh2* | -1.69 | 9.23E-10 |
|  |  |  | Gohir.D10G160800 | *AMT1;1* | -2.32 | 2.40E-13 |
|  |  |  | Gohir.D10G160900 | *AMT1;1* | -2.27 | 9.53E-13 |
|  |  |  | Gohir.A10G106700 | *AMT1;1* | -4.10 | 4.13E-13 |

|  |  |  |  |  |
| --- | --- | --- | --- | --- |
| bb | Gohir.D02G050000 | *AMT1;1* | -1.15 | 3.78E-12 |
|  | Gohir.A10G106400 | *AMT1;1* | -2.87 | 1.40E-12 |
|  | Gohir.D10G160800 | *AMT1;1* | -1.59 | 1.34E-13 |
|  | Gohir.A10G106700 | *AMT1;1* | -1.38 | 2.78E-07 |
| soa | Gohir.A10G106400 | *AMT1;1* | -2.14 | 3.13E-14 |
| sob | Gohir.A03G061200 | *gdh2* | -0.80 | 8.43E-14 |
|  | Gohir.A02G044000 | *AMT1;1* | -1.13 | 5.48E-15 |
|  | Gohir.A10G106400 | *AMT1;1* | -6.61 | 5.50E-13 |
|  | Gohir.D10G160800 | *AMT1;1* | -2.63 | 5.17E-13 |
|  | Gohir.A10G106700 | *AMT1;1* | -4.48 | 2.77E-14 |
| boa | Gohir.D11G140500 | *AMT1;1* | -5.5 | 1.50E-05 |
|  | Gohir.D10G160800 | *AMT1;1* | -1.86 | 2.76E-14 |
|  | Gohir.A10G106400 | *AMT1;1* | -3.37 | 8.51E-13 |
|  | Gohir.A10G106700 | *AMT1;1* | -8.35 | 6.18E-14 |

a Arabidopsis and *Gossypium hirsutum* genes identification.
b log fold change (log 2FC) between two replicated experiments.
c Transcripts exhibiting a FDR lower than 0.0001%.

### 3.5 Effects of nitrogen reduction combined with biochar on seedling growth

Biochar combined with different urea significantly reduced the active absorption area of roots, with bbf treatment <0.1mm reducing the root area by 19.31%, and boaf treatment with fine roots reducing the active absorption area by 11.30%.The active absorption area of fine roots treated by soaf increased by 9.43 cm2. The active absorption area of 0.1~2mm roots increased by 1.80~9.10 cm2 (Figure 4A).The change rule of root fresh weight was similar to that of active absorption area, that is, fresh weight of <0.1mm fine roots decreased, and that of 0.1-2mm roots slightly increased, among which fresh weight of SBF and boaf treated fine roots significantly decreased (*P*<0.05), fresh weight of roots 0.1~2mm treated by BBM significantly decreased (*P*<0.05), and fresh weight of roots soam significantly increased (*P*<0.01)(Figure. 4B). The nitrogen fertilizer agronomic efficiency of soa treatment increased by 1.49 kg kg^-1^, whereas that of boa treatment decreased by 0.91 kg kg^-1^. The remaining treatments showed an increasing trend (Figure 4C).

**Figure 4.**
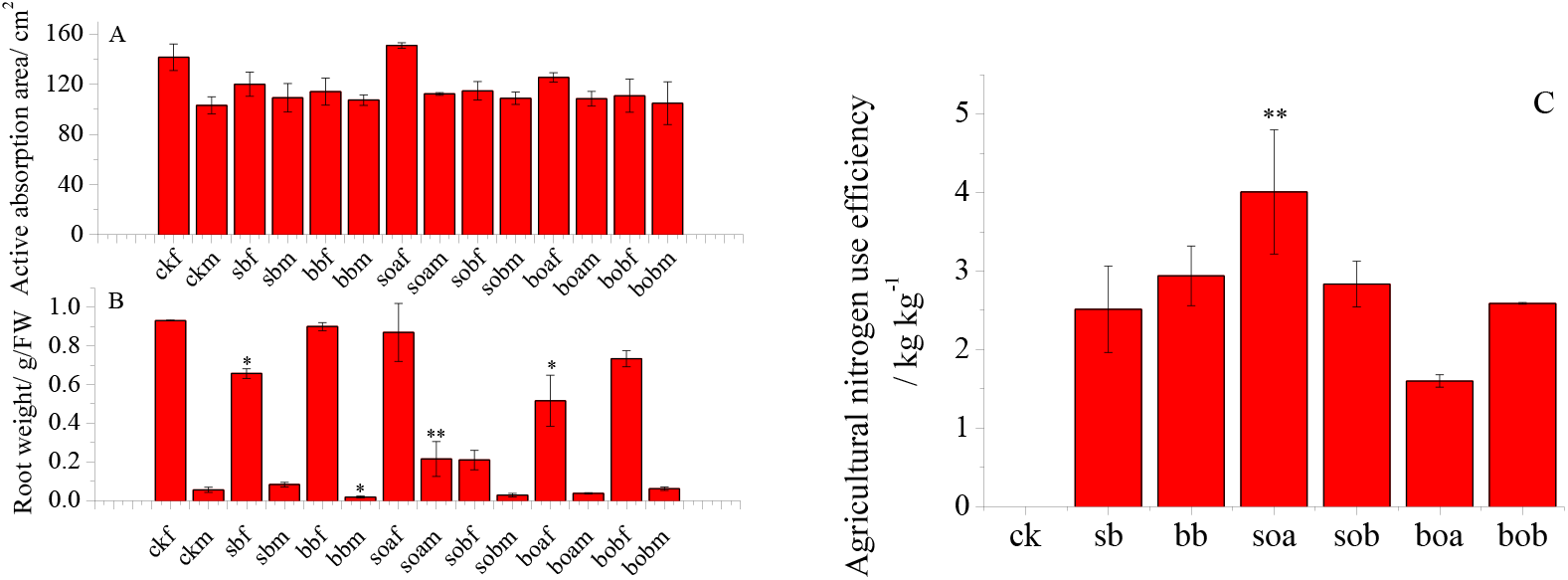
Differences in root active absorption area, fresh weight and seedling nitrogen agricultural efficiency at the classification of root between *d*<0.1mm and 0.1<*d*<2mm. Results are presented as mean values for seven plants with SE. Asterisk indicate significant differences with a confidence interval at \**P* < 0.05 and \*\**P*<0.01, respectively.

## 4 Discussions

Improving nitrogen use efficiency of crops can not only reduce planting costs, but also reduce energy consumption caused by chemical fertilizers, and fundamentally alleviate global climate change (Sinha et al., 2015; Zuluaga et al. 2017). Regulation of root system (morphology, configuration, physiology, molecular level, etc.) is one of the key measures to achieve the above goals (Kumar et al. 2018). Thus, regulation of physiological characteristics can increase the affinity of crop roots to NH_4_^+^ and improve nitrogen utilization efficiency, the core of which is to improve the activity of key metabolic enzymes of nitrogen (Perchlik & Tegeder 2017). Generally speaking, the activity of NR, GDH and GS decreased with the decrease of nitrogen, while GOGAT was associated with nitrogen phenotype, that is, the activity was still high under low nitrogen level, for instance, there are some data implying that intraspecific root variation (Liu et al. 2018) and changes in rhizosphere nitrogen availability and biochemical processes improved nitrogen efficiency and thus promoted crop growth (Billah et al. 2019). Our results are similar to above conclusions, but we considered that the causes should include at least the effects of microbial changes in the fine root rhizosphere, especially the increased number of ammonia-oxidizing bacteria.

Of course, main nitrogen source (NH_4_^+^ or NO_3_^-^) induced by ammonia-oxidizing bacteria through biochar, hereinto, NH_4_^+^ is higher plants used for synthesis of amino acids and nucleic acid, it into the cells of the flux is mainly composed of ammonium transporters (ammonium transporters, *AMTs*) regulation (Kumar et al. 2003). *AMTs* in plants are encoded by *AMT1* and *AMT2* subfamilies, among which *AMT1s* is associated with efficient NH_4_^+^ transport in plants. *AMT1;1* in *Arabidopsis thaliana* is mainly expressed in roots and leaves, *AMT1;3* is only expressed in the root, and *AMT1;1* and *AMT1;3* is sensitive to nitrogen deficiency (Wirén 2000). When nitrogen is insufficient, *AMT1;1* and *AMT1;3* was significantly up-regulation in root dermis including root hairs and *AMT1.3* May be associated with C and N metabolism in roots (Dominique et al. 2006).This study showed that *AMT1;2* was significantly up-regulated in diameter <0.1mm roots, and decreased to varying degrees in roots at class between 0.1 and 2mm. It suggests that biochar could more easily induce <0.1mm root *AMT1;1* change.

Glutamate dehydrogenase (GDH) is a rate-limiting enzyme for plants to complete Ammonium assimilation under biological or abiotic stress, in which the metabolic process involving NADH-*gdh* mainly occurs in the root system (Andrews et al. 2004; Asif et al. 2020). If the environment is dark or carbon stressed, a reversible reaction will occur to provide carbon framework for the tricarboxylic acid cycle (Kishorekumar et al. 2020; Fontaine et al. 2012), indicating that the activity of GDH, an intermediate involved in carbon and nitrogen metabolism, is closely related to the environment. This study showed that 20% and 50% reduction of nitrogen, GDH activity in fine root increased significantly, but its increase in medium root is not obvious or significantly reduced, because the biological carbon and nitrogen under the condition of reduction, the formation of pH of 9.8 degrees and nitrogen limited in alkaline environment, stimulated the GDH protein coding, improved GDH activity, and biochar enhanced peanut seedling photosynthesis (Xu et al. 2015), make no carbon inhibitory effect of seedling or strength is weak. On the other hand, biochar increased the content of soil organic matter, which may make up for the lack of carbon to some extent. Moreover, biochar produces a relatively dark environment, which may make the proportion of C: N in the root system more reasonable (Chen et al. 2014). This was also consistent with a number of studies (Farhangi-abriz & Torabian 2018; Noguera et al. 2012). Generally, higher plant GDH has a weak affinity for NH_4_^+^, so NH_4_^+^ is absorbed mainly by glutamine synthase (GS)/ glutamic synthase (GOGAT) pathway (Chen et al., 2019). The results of this experiment showed that, under the action of biochar and nitrogen reduction, the GOGAT activity of seedling fine roots was strongly inhibited, while the interference of medium roots was weak but increased. This indicates that during the process of ammonium assimilation in the induced GS/GOGAT pathway, exogenous ammonium ions did not decrease and even increased to some extent, because biochar increased the microbial diversity of medium rhizosphere ammonia-oxidizing microorganisms, leading to a significant increase in rhizosphere nitrate ions.

Nitrate reductase (NR) was one of the key enzymes of nitrogen metabolism in high plants, and NO_3_^-^ and light had a strong influence on it (Nemie-Feyissa et al., 2013; Chow et al., 2004). The results showed that the activity of NR treated with sobm increased by 83%, and the activity of NR treated with sobm increased slightly, while the activity of NR treated with other fine roots decreased. This suggested that NO_3_^-^ level contributes less to cotton seedling metabolism, and NR activity may be related to root senescence under the biochar-protection. Feng et al. (2019) have speculated that hormone substances contained in biochar may promote the improvement of NR activity. Researchers have shown that hormone gene has an evident in nitrogen metabolism (Zuluaga et al. 2017). In their opinions, selection of plants, from either a genetically manipulated population or genetic resources, with expression of nitrate reductase/nitrite reductase primarily in the root or shoot should increase plant/crop growth and hence yield under specific environmental conditions where a large number of experiments confirmed that biochar significantly changed the rhizospheric character (Olmo et al. 2014;; Farhangi-abriz & Torabian 2017). In dark conditions, fine roots produced more cytokinin and hence increase nitrate reductase activity, while biochar prolongs darkness and promotes nitrogen assimilation (Figure 3A).

In short, we detected that when urea dose reduction by 20% with biochar was applied, the promoting effect increased by significantly up-regulated the expression levels of *AMT1;1* and *gdh2* which improved ammonium assimilation, and it was also closely related to the crosstalk between *AMT* and *gdh3*. And the stimulation of cytokinin to nitrate reductase activity in fine root increased the substrate concentration of ammonium assimilation.

## 5 Conclusions

Results presented here indicate that crosstalk between *gdh2* and *AMT1;1* signaling pathway coordinates ammonia assimilation of *Gossypium hirsutum* under biochar and limited urea condition which exhibits a 1.51 kg kg^-1^ to 3.99 kg kg^-1^ higher nitrogen agronomic efficiency than the control at the seedling stage. It is tempting to speculate this difference is the direct result of *gdh2* up-regulation and *gdh3* supplement. GOGAT, NR, GPT and GDH enzyme activities all improvement could account for its higher nitrogen assimilation induced by biochar if the root nitrogen assimilation via GS/GOGAT and GDH pathway is definitely dependent upon the supplement of *gdh3* expression where is related to the function of the root system.

We further found that cytokinins both activated and inhibited nitrate reductase activity under biochar condition, which is directly related to root turnover rate or the age and function of the root. That is, if the target is a fine root (*d*<0.1mm), it is shown as excitation between both cytokinins and nitrate reductase, and if a medium root (0.1mm<*d*<2mm), it is shown as antagonism between the two enzymes. Expect for, biochar location seems partial authentic as an explanation for these data.

Overall, biochar induced *gdh3* expression promotion GS/GOGAT and GDH pathway might account for improving nitrogen assimilation in fine root.

## Abbreviation

AOB: rhizospheric ammonia-oxidizing bacteria;
NR: nitrate reductase;
GPT: glutamic-pyruvic transaminase activity;
GDH: glutamate dehydrogenase;
GOGAT: Glutamine oxoglutarate amino transferase

## Acknowledgements

This work was supported by the National Natural Science Foundation of China, with a grant to R.L. (31660073). We thank Xu CY for her suggestions on the first version of the manuscript and Zhang YS, Jin YY and Bai DW for support with part of the analysis. We also thank the Institute of Soil Fertilizer and Agricultural Water Conservation of Xinjiang Academy of Agricultural Sciences for their support and staff assistance during the fieldwork.

## References

Andrews, M., Lea, P. J., Raven, J. A., & Lindsey, K.. (2004). Can genetic manipulation of plant nitrogen assimilation enzymes result in increased crop yield and greater n - use efficiency? an assessment. Annals of Applied Biology, 145(1), 25–40. https://doi.org/10.1111/j.1744-7348.2004.tb00356.x.

Asif I., Dong Q., Wang Z., Wang X. R., Gui H.P., Zhang H. H., Pang N.C., Zhang X. L. and Song M. Z.. (2020). Growth and nitrogen metabolism are associated with nitrogen-use efficiency in cotton genotypes, Plant Physiology and Biochemistry, 149, 61–74. DOI: 10.1016/j.plaphy.2020.02.002.

Backer, R. G. M., Saeed, W., Seguin, P., & Smith, D. L.. (2017). Root traits and nitrogen fertilizer recovery efficiency of corn grown in biochar-amended soil under greenhouse conditions. Plant and Soil, 415(1-2), 465–477. DOI 10.1007/s11104-017-3180-6.

Bian R. J., Joseph S., Wei, & Shi, et al. (2019). Biochar DOM for plant promotion but not residual biochar for metal immobilization depended on pyrolysis temperature. The Science of the total environment. DOI: 10.1016/j.scitotenv.2019.01.224.

Billah, M. M., Ahmad, W., & Ali, M.. (2019). Biochar particle size and rhizobia strains effect on the uptake and efficiency of nitrogen in lentils. Journal of Plant Nutrition, 42(2), 1–17. DOI: 10.1080/01904167.2019.1628984.

Chen, C. P., Cheng, C. H., Huang, Y. H., Chen, C. T., Lai, C. M., & Menyailo, O. V., et al. (2014). Converting leguminous green manure into biochar: changes in chemical composition and C and N mineralization. Geoderma, 232-234, 581–588. DOI: 10.1016/j.geoderma.2014.06.021.

Chen M. D., Liu C., Liu D. R., Tang Q. L., Lin J. Z., & Liu X. M.. (2019). Ectopic Expression of a Fungal AbGDH Gene from Alternaria brassicicola Improves Nitrogen Use Efficiency in Rice. Life Science Research, 23(1): 1–12. DOI: CNKI: SUN:SMKY.0.2019-01-001.

Chow, F., Oliveira, M. C. D., & Marianne Pedersén. (2004). In vitro assay and light regulation of nitrate reductase in red alga gracilaria chilensis. Journal of Plant Physiology, 161(7), 0–776. DOI: 10.1016/j.jplph.2004.01.002.

Dilfuza, E., Stephan, W., Undine, B., Abd_Allah, E. F., & Gabriele, B.. (2016). Biochar treatment resulted in a combined effect on soybean growth promotion and a shift in plant growth promoting rhizobacteria. Frontiers in Microbiology, 7. DOI: 10.3389/fmicb.2016.00209.

Dominique Loqué, Yuan, L., Kojima, S., Gojon, A., Wirth, J., & Gazzarrini, S., et al. (2006). Additive contribution of amt1;1 and amt1;3 to high-affinity ammonium uptake across the plasma membrane of nitrogen-deficient *Arabidopsis* roots. Plant Journal, 48(4), 522–534. DOI: 10.1111/j.1365-313x.2006.02887.x.

Farhangi-Abriz, S., & Torabian, S.. (2017). Biochar increased plant growth-promoting hormones and helped to alleviates salt stress in common bean seedlings. Journal of Plant Growth Regulation, 37(2), 591–601. DOI:10.1007/s00344-017-9756-9.

Farhangi-Abriz, S., & Torabian, S.. (2018). Biochar improved nodulation and nitrogen metabolism of soybean under salt stress. Symbiosis, 74(3), 1–9. DOI: 10.1007/s13199-017-0509-0.

Feng, L., Xu, W. L., Tang, G. M., Sun, N. C., Pu, S. H., Geng, Z. C.. (2019). Effects of Biochar Combined with Nitrogen on Root Morphology and System Architecture during *Gossypium hirsutum* L. full bloom Stage. Transactions of the Chinese Society for Agricultural Machinery, 50(3):241–249. DOI: 10.6041/j.issn.1000-1298.2019.03.026

Fontaine, J. X., Terce-Laforgue, T., Armengaud, P., Clement, G., Renou, J. P., & Pelletier, S., et al. (2012). Characterization of a NADH-dependent glutamate dehydrogenase mutant of *Arabidopsis* demonstrates the key role of this enzyme in root carbon and nitrogen metabolism. The Plant Cell, 24(10), 4044–4065. DOI: 10.1105/tpc.112.103689

French, E., & Iyer-Pascuzzi, A. S.. (2018). A role for the gibberellin pathway in biochar-mediated growth promotion. Scientific Reports, 8(1), 5389. DOI: 10.1038/s41598-018-23677-9.

Hale, L., Luth, M., & Crowley, D.. (2015). Biochar characteristics relate to its utility as an alternative soil inoculum carrier to peat and vermiculite. Soil Biology and Biochemistry, 81(81), 228–235. DOI: 10.1016/j.soilbio.2014.11.023.

Hashem, A., Kumar, A., Abeer, & M, et al. (2019). Arbuscular mycorrhizal fungi and biochar improves drought tolerance in chickpea. Saudi journal of biological sciences. https://doi.org/10.1016/j.sjbs.2018.11.005.

Hong, F., Breitling, R., Mcentee, C. W., Wittner, B. S., Nemhauser, J. L., & Chory, J.. (2006). Rankprod: a bioconductor package for detecting differentially expressed genes in meta-analysis. Bioinformatics, 22(22), 2825–2827. DOI: 10.1093/bioinformatics/btl476.

Jeffery, S., Verheijen, F. G. A., Velde, M. V. D., & Bastos, A. C.. (2011). A quantitative review of the effects of biochar application to soils on crop productivity using meta-analysis. Agriculture Ecosystems & Environment, 144(1), 175–187. DOI: 10.1016/j.agee.2011.08.015.

Kishorekumar R., Bulle M., Wany A., Gupta K.J. (2020) An Overview of Important Enzymes Involved in Nitrogen Assimilation of Plants. In: Gupta K. (eds) Nitrogen Metabolism in Plants. Methods in Molecular Biology, 2057. Humana, New York, NY. https://xs.scihub.ltd/ https://doi.org/10.1007/978-1-4939-9790-9_1.

Kolton, M., Graber, E. R., Tsehansky, L., Elad, Y., & Cytryn, E.. (2017). Biochar-stimulated plant performance is strongly linked to microbial diversity and metabolic potential in the rhizosphere. New Phytologist, 213(3), 1–12. https://doi.org/10.1111/nph.14253.

Konishi, N., Ishiyama, K., Matsuoka, K., Maru, I., Hayakawa, T., & Yamaya, T., et al. (2014). Nadh-dependent glutamate synthase plays a crucial role in assimilating ammonium in the arabidopsis root. Physiologia Plantarum, 9(8), 138–151. DOI: info:doi/10.1111/ppl.12177.

Kumar, A., Silim, S. N., Okamoto, M., Siddiqi, M. Y., & Glass, A. D. M.. (2003). Differential expression of three members of the amt1 gene family encoding putative high-affinity NH_4_^+^ transporters in roots of oryza sativa subspecies indica. Plant, Cell & Environment, 26. DOI:10.1046/j.1365-3040.2003.01023.x.

Kumar, S. S., Akanksha, T., & Kumar, M. P.. (2018). External nitrogen and carbon source-mediated response on modulation of root system architecture and nitrate uptake in wheat seedlings. Journal of Plant Growth Regulation. DOI: 10.1007/s00344-018-9840-9.

Li X. L., Zheng X. F., Jiang Y. M., & Wei S. C.. (2019). Effect of ethephon on absorption, distribution and utilization of nitrate-nitrogen by T337 apple dwarf rootstock seedlings. Plant Nutrition and Fertilizer Science, 25(7), 1204–1210.

Liu F. C., Ma H. L., Du Z.Y., Ma B. Y., Jing D. W., 2018. Response of endogenous hormones and key enzyme of nitrogen metabolism among different branch orders of fine root in the incision of poplar to root pruning. Ecology and Environmental Sciences, 27(12): 2234–2242. DOI: 10.16258/j.cnki.1674-5906.2018.12.008.

Lonardo, S. D., Vaccari, F. P., Baronti, S., Capuana, M., Bacci, L., & Sabatini, F., et al. (2013). Biochar successfully replaces activated charcoal for in vitro culture of two white poplar clones reducing ethylene concentration. Plant Growth Regulation, 69(1), 43–50. DOI: 10.1007/s10725-012-9745-8.

Lu, Y., Mengjie, Y., Xing, L., Caixian, T., Xingmei, L., & Brookes, P. C., et al. (2018). Combined application of biochar and nitrogen fertilizer benefits nitrogen retention in the rhizosphere of soybean by increasing microbial biomass but not altering microbial community structure. Science of the Total Environment, 640-641, 1221–1230. DOI: 10.1016/j.scitotenv.2018.06.018.

Masiello, C. A., Chen, Y., Gao, X., Liu, S., & Silberg, J. J.. (2013). Biochar and microbial signaling: production conditions determine effects on microbial communication. Environmental Science & Technology, 47(20), 11496–11503. DOI: 10.1021/es401458s.

Mehari, Z. H., Elad, Y., Rav-David, D., Graber, E. R., Meller H. Y..(2015). Induced systemic resistance in tomato (*Solanum lycopersicum*) againstbotrytis cinereaby biochar amendment involves jasmonic acid signaling. Plant and Soil, 395(1-2), 31–44. DOI: 10.1007/s11104-015-2445-1.

Nacry, P., Eléonore Bouguyon, & Gojon, A.. (2013). Nitrogen acquisition by roots: physiological and developmental mechanisms ensuring plant adaptation to a fluctuating resource. Plant and Soil, 370(1-2), 1–29. DOI: 10.1007/s11104-013-1645-9.

Nemie-Feyissa, D., Królicka, Adriana, F?Rland, N., Hansen, M., Heidari, B., & Lillo, C.. (2013). Post-translational control of nitrate reductase activity responding to light and photosynthesis evolved already in the early vascular plants. Journal of Plant Physiology, 170(7), 662–667. DOI: 10.1016/j.jplph.2012.12.010.

Noguera, D., Barot, S., Laossi, K. R., Cardoso, J., Lavelle, P., & Carvalho, M. H. C. D.. (2012). Biochar but not earthworms enhance rice growth through increased protein turnover. Soil Biology & Biochemistry, 52, 13–20. DOI: 10.1016/j.soilbio.2012.04.004.

N von Wirén. (2000). The molecular physiology of ammonium uptake and retrieval. Current Opinion in Plant Biology, 3(3), 254–261. DOI: 10.1016/S1369-5266(00)00073-X.

Olmo, M., Alburquerque, José Antonio, Barrón, Vidal, del Campillo, María Carmen, Gallardo, A., & Fuentes, M., et al. (2014). Wheat growth and yield responses to biochar addition under mediterranean climate conditions. Biology and Fertility of Soils, 50(8), 1177–1187. DOI:10.1007/s00374-014-0959-y.

Pandey, V., Patel, A., & Patra, D. D.. (2016). Biochar ameliorates crop productivity, soil fertility, essential oil yield and aroma profiling in basil (*Ocimum basilicum* l.). Ecological Engineering, 90, 361–366. DOI: 10.1016/j.ecoleng.2016.01.020.

Perchlik, M., & Tegeder, M.. (2017). Improving plant nitrogen use efficiency through alteration of amino acid transport processes. Plant physiology, 175(1), pp.00608.2017. DOI: 10.1104/pp.17.00608.

Polzella, A., De Zio, E., Arena, S., Scippa, G. S., Scaloni, A., & Montagnoli, A., et al. (2018). Toward an understanding of mechanisms regulating plant response to biochar application. Plant Biosystems / An International Journal Dealing with all Aspects of Plant Biology, 1–10. DOI: 10.1080/11263504.2018.1527794.

Razaq, M., Salahuddin., Shen, H. L., Sher, H., & Zhang, P.. (2017). Influence of biochar and nitrogen on fine root morphology, physiology, and chemistry of acer mono. Scientific Reports, 7(1), 5367. DOI: 10.1038/s41598-017-05721-2.

Sarasin, A., & Kauffmann, A. (2008). Overexpression of DNA repair genes is associated with metastasis: a new hypothesis., 659(1), 49–55. DOI:10.1016/j.mrrev.2007.12.002.

Sinha S.K., Rani M., Bansal N., Gayatri Venkatesh K., & Mandal P.K.. (2015). Nitrate starvation induced changes in root system architecture, carbon:nitrogen metabolism, and mirna expression in nitrogen-responsive wheat genotypes. Applied Biochemistry and Biotechnology, 177(6), 1299–1312. DOI: 10.1007/s12010-015-1815-8.

Song, Y., Zhang, X., Ma, B., Chang, S. X., & Gong, J.. (2014). Biochar addition affected the dynamics of ammonia oxidizers and nitrification in microcosms of a coastal alkaline soil. Biology & Fertility of Soils, 50(2), 321–332. DOI: 10.1007/s00374-013-0857-8.

Spokas, K. A., Baker, J. M., & Reicosky, D. C.. (2010). Ethylene: potential key for biochar amendment impacts. Plant and Soil, 333(1-2), 443–452. DOI: 10.1007/s11104-010-0359-5.

Xu, C. Y., Hosseini-Bai, S., Hao, Y., Rachaputi, R. C. N., Wang, H., & Xu, Z., et al. (2015). Effect of biochar amendment on yield and photosynthesis of peanut on two types of soils. Environmental Science and Pollution Research, 22(8), 6112–6125. DOI: 10.1007/s11356-014-3820-9.

Turano, F. J. (1996). Purification of mitochondrial dehydrogenase from dark-grown soybean seedlings. Plant Physiology, 112(3), 337–344. DOI: 10.1034/j.1399-3054.1998.1040307.x.

Viger, Maud, Miglietta, Franco, Taylor, & Gail, et al. (2015). More plant growth but less plant defence? first global gene expression data for plants grown in soil amended with biochar. Global change biology bioenergy, 7, 658–672. https://doi.org/10.1111/gcbb.12182.

Xu Z. Z., & Zhou G. S.. (2004). Research advance in nitrogen metabolism of plant and its environmental regulation. Chinese Journal of Applied Ecology, 15(3), 511–516. DOI: http://dx.doi.org/.

Waqas, M., Khan, A. L., Kang, S.-M., Kim, Y.-H., & Lee, I.-J. (2014). Phytohormone-producing fungal endophytes and hardwood-derived biochar interact to ameliorate heavy metal stress in soybeans. Biology and Fertility of Soils, 50(7), 1155–1167. DOI:10.1007/s00374-014-0937-4.

Wu, H., Zeng, G., Liang, J., Chen, J., Xu, J., & Dai, J., et al. (2016). Responses of bacterial community and functional marker genes of nitrogen cycling to biochar, compost and combined amendments in soil. Applied Microbiology and Biotechnology, 100(19), 8583–8591. DOI:10.1007/s00253-016-7614-5.

Zuluaga, D. L., Domenico, D. P., Michela, J., Luca, C. P., Gabriella, S., & Turgay, U.. (2017). Durum wheat mirnas in response to nitrogen starvation at the grain filling stage. PLOS ONE, 12(8), e0183253-.

